# Attenuation of colitis-induced visceral hypersensitivity and pain by silencing TRPV1-expressing fibers in rat colon

**DOI:** 10.1101/2023.06.27.546676

**Authors:** Yoav Mazor, Nurit Engelmayer, Halla Nashashibi, Lisa Rottenfußer, Shaya Lev, Alexander M Binshtok

## Abstract

**Background and Aims:** Abdominal pain in patients with inflammatory bowel disease (IBD) is common and debilitating. In our study, we aim to utilize transient receptor potential vanilloid 1 (TRPV1) channels, large-pore cation channels expressed on nociceptors, as a drug delivery system to selectively inhibit visceral nociceptors and thus visceral pain in a rodent model of IBD.

**Methods:** We induced colitis in rats using intrarectal dinitrobenzene sulfonic acid. Visceral hypersensitivity, spontaneous pain, and responsiveness of the hind paws to noxious heat stimuli were examined before and after the intrarectal application of sodium channel blocker QX-314 alone or together with TRPV1 channel activators or blockers.

**Results:** Intrarectal co-application of QX-314 with TRPV1 channel activator capsaicin significantly inhibited colitis-induced gut hypersensitivity. Furthermore, in the model of colitis, but not in naïve rats, QX-314 alone was sufficient to reverse gut hypersensitivity. The blockade of TRPV1 channels prevented this effect of QX-314. Finally, applying QX-314 alone to the inflamed gut inhibited colitis-induced ongoing pain.

**Conclusions:** Selective silencing of nociceptors by QX-314 entering via exogenously or endogenously activated TRPV1 channels diminish IBD-induced gut hypersensitivity. These results yet again confirm the central role of TRPV1-expressing nociceptive neurons in IBD pain. The lack of QX-314 effect on naïve rats suggests its selective analgesic effect in IBD pain. Moreover, our results demonstrating the effect of QX-314 alone imply the role of a tonically active TRPV1 channel in the pathophysiology of IBD pain. This approach provides proof-of-concept for using charged activity blockers for selective and effective blockade of visceral pain.

## Introduction

Abdominal pain is the presenting symptom in up to 70% of patients with inflammatory bowel disease (IBD), and at least one-third of patients have persistent pain despite optimal anti-inflammatory treatment^1^. IBD-induced pain is associated with higher anxiety levels, depression, social dysfunction, and work disability, thus devastatingly affecting patients’ lives^2–4^. Available analgetic treatments are non-specific, with variable effectiveness and considerable side effects, including addiction and increased morbidity^5–7^.

Recent studies have made substantial progress in understanding gut innervation by nociceptive neurons and proposed potential molecular and cellular mechanisms underlying IBD-induced pain^8–11^. Several lines of evidence demonstrate the role of the transient receptor potential V1 channel (TRPV1) in IBD-induced pain in human and animal models of colitis^11–18^. This large pore cation non-selective transducer channel is expressed by both somatic nociceptive neurons and nociceptive peptidergic neurons innervating the gut^13^. The activation of these channels by various proinflammatory mediators contributes to inflammatory pain^19–22^. In animal models of IBD, systemic blockade of TRPV1 reduces the intensity of visceral hypersensitivity, implying that abnormal activity of TRPV1 contributes to IBD-induced visceral pain^12^. Nevertheless, selective blockers of TRPV1 have failed in human studies due to life-threatening side effects, specifically hyperthermia^23^.

In this study, we aimed to study the effect of selective blockade of TRPV1-expressing nociceptive fibers in IBD-induced pain. We utilized a selective silencing approach we have previously developed to produce pain-selective anesthesia and examine the role of TRPV1-expressing fibers in somatic pain and itch^24, 25^. In this approach, we used otherwise membrane impermeant sodium channel blocker QX-314 and shuttled it into nociceptive neurons through TRPV1 channels activated by capsaicin. We showed that the co-application of capsaicin and QX-314 led to selective blockade of nociceptive activity and inhibited pain without affecting other sensory or motor modalities^24^. Furthermore, we and others showed that tonically open TRPV1 channels, following endogenous activation by either pruritogens or proinflammatory mediators, were sufficient to allow entry of QX-314 into neurons, allowing for dissecting the role of these neurons in pain and itch^25, 26^.

Here, we show that in dinitrobenzene sulfonic acid (DNBS)-induced colitis in rats, silencing TRPV1-expressing neurons by intrarectal co-application of capsaicin with QX-314 reverses gut hypersensitivity. These results suggest that TRPV1-expressing nociceptive neurons innervating the gut play a role in colitis-induced pain. We further show that intrarectal administration of QX-314 alone in colitis conditions was sufficient to abolish colitis-induced visceral hypersensitivity and ongoing pain. Notably, the application of QX-314 alone in naïve animals did not affect the normal gut sensitivity, suggesting the selective effect of QX-314 on colitis-induced pain. These results propose that in an IBD model, TRPV1 channels are tonically active, allowing entry of QX-314 into nociceptive fibers innervating the gut. Indeed, the pharmacological blockade of TRPV1 channels prevented the effect of QX-314 alone on colitis-induced gut hypersensitivity. Our results demonstrate that TRPV1 channels activated during IBD may serve as a “natural” drug delivery system, laying the foundation for developing this platform as an effective and selective treatment for IBD-induced abdominal pain.

## Materials and Methods

### Animals

All of the procedures performed in this study were in full accordance with the guidelines of the Hebrew University Animal Care Committee and were approved and ratified by the Committee. Procedures were performed on male and female adult (250-274 g) Sprague-Dawley rats. The rats were housed in groups of four under a 12-h light/dark cycle. Rats were habituated to the testing environment and handlers prior to procedures. The room temperature and humidity remained stable during the entire experiment, and food and water were regularly provided. Prior to all experiments described, the animals went through a sufficient period of habituation to the testing apparatus used and the experimental environment.

### Colitis induction

Following acclimation to the animal facility, animals were anesthetized in an isoflurane chamber with a continuous flow of 4% isoflurane. Once sedated, a nose cone with a continuous flow of 1-2% isoflurane was applied. The animals’ hind legs were taped to the surface of a plastic container and placed at a 45^0^ angle by raising the container edge. 30 mg in 250 μl of DNBS solution was administered transanally using a flexible tube inserted 6 cm proximal to the anal canal. The animals were left at a 45^0^ angle for 90 seconds to avoid excretion of the DNBS solution. Animals were then allowed to recover and monitored twice daily for 3 days to assess for clinical signs of inflammation and weight loss. All experiments were conducted on day 4 after colitis induction, following which the animals were humanely sacrificed and the ano-colon carefully dissected, removed, and examined to ensure colitis induction (**Supplementary Figure 1**).

### Intrarectal Drug Delivery

All treatments were administered via enema in isoflurane-sedated animals, as described above. During administration, the animals were placed at a 45^0^ angle for 90 seconds to reduce the excretion of the drug/vehicle. Animals were allowed to recover for 10 minutes after awakening from anesthesia until fully mobile.

### Drugs

Dinitrobenzene sulfonic acid (DNBS, Sigma-Aldrich, 556971) was prepared as follows: 240 mg of DNBS was dissolved in 1 ml of 100% ethanol and 1 ml of DDW, and each animal was administered 250 μl (30 mg) of the solution. Capsaicin (Sigma-Aldrich, 21750) was prepared by dissolving 5 mg of capsaicin in 0.5 ml TWEEN (Sigma-Aldrich, P9416), 0.5 ml ethanol, and 4 ml DDW, and each animal was administered 1 ml of solution. 2% lidocaine and 2% QX-314 (N-ethyl-lidocaine) solutions were prepared by dissolving 80 mg of either lidocaine or QX-314 in 4 ml DDW, and each animal was administered 1 ml of solution. Capsazepine (Sigma-Aldrich, P9416) was prepared by dissolving either 5, 10, 12.5, or 15 mg of capsazepine in 2.5 ml TWEEN (Sigma-Aldrich P9416) and 2.5 ml ethanol, and each aminal received 1 ml of solution. The vehicle was prepared using the same solvent solution used for the active components in the experimental groups without the addition of the active component.

### Assessment of Visceral Hypersensitivity

Colorectal distention by inflatable balloon was used to assess visceral hypersensitivity^27^. Three radial holes were carved into the terminal end of a flexible tube. Then, a 6 cm balloon was constructed from a piece of latex (Durex) and was tied to the end of the flexible tube using dental floss (Dr. Collins). The other end of the tube was connected to a barometer with a tri-directional valve, and a 10 ml syringe was used to manually fill the balloon and maintain the required pressure. The balloon lacked inherent compliance at volumes of up to 10 ml, used in all experiments.

Animals were anesthetized, and the balloon was lubricated (Sion Biotext) and inserted into the distal colon, 0.5 cm from the anus. Fabric tape (Life) was used to secure the balloon to the tail, and the animals were allowed to recover for 10 minutes until fully mobile. After that, the balloon was inflated to a pressure of either 40 mmHg for measuring capsaicin– or colitis-induced hypersensitivity or 60 mmHg in naïve rats (**Figure 1A**). The pressure was kept constant over the duration of 8 minutes, and abdominal withdrawal reflexes (AWRs) were visually counted and used as a measure of visceral hypersensitivity.

**Figure 1.**
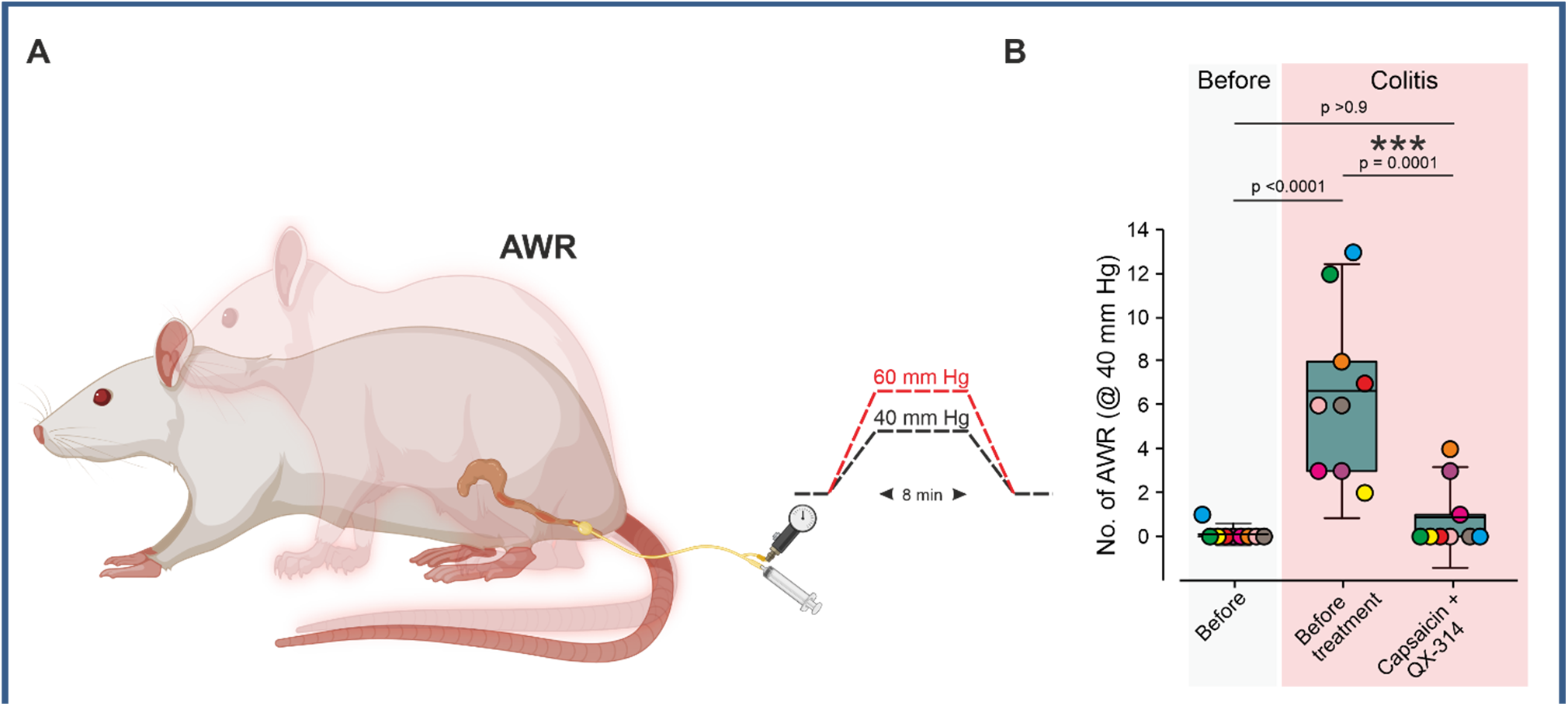
Co-application of capsaicin and QX-314 attenuates gut hypersensitivity induced by DNBS, in a rodent model of colitis. A. The experimental setup. A colorectal balloon connected to a barometer is inserted into the rat’s colon. Abdominal withdrawal reflexes (AWRs) are defined as an enhanced back kyphosis and neck extension and are measured following colorectal distension resulting from applying either 40 mm Hg (colitis model) or 60 mm Hg (naïve rats). **B.** Box plot and individual values of the number of AWRs following colorectal distension at 40 mm Hg in the same rats (color-coded) before and 4 days after treatment with DNBS (*colitis*) and 4 days after DNBS but treated with an intrarectal co-application of 1 mg/ml capsaicin and 20 mg/ml QX-314 (2%). Note that treatment with DNBS significantly increases gut sensitivity, which was reversed to control values by treatment with capsaicin and QX-314. N=9, RM One-way ANOVA with post hoc Bonferroni. The box plot depicts the mean, 25-75% percentile, and the whiskers depict 1.5 SD.

AWRs were defined as characteristic involuntary responses that involve sudden kyphosis, cervical extension, and occasional vocalization^28^. AWRs were quantified visually during the experiment, and all experiments were filmed and recorded for post hoc data validation.

### Assessment of noxious thermal threshold

The response to noxious pain stimuli was assessed using the Hargreaves heat latency test (Ugo-Basile, Varese, Italy). The rats were placed on a plexiglass surface, and a laser beam calibrated to heat the paw to 52^0^ was directed towards the plantar aspect of the hind paw. Paw withdrawal latency (PWL), the time to a behavioral withdrawal response, was measured^29, 30^.

### Assessment of ongoing pain, pain grimace scale

To test ongoing pain behavior without evocation, facial characteristic analysis was performed prior and following inflammation with and without treatments. Rats were habituated to room and chambers for 3 days prior to experiments. Afterward, animals were placed in a plexiglass chamber, and headshots were taken randomly 3 times over 20 minutes prior to inducing DNBS colitis. 4 days after inducing colitis, animals were placed in the plexiglass chambers and imaged randomly 3 times over 20 minutes. Then, each animal was given a dose of QX-314 (20mg/ml) and left to rest for 10 minutes. Following the treatment, 3 random images were recorded over the span of 20 minutes. Images were coded by blinded testers, and photos were randomized and scored (with an intensity rate of 0, 1 or 2) by the following criteria: orbital tightening, nose/cheek bulge, ear position and whisker change. In each case, 0 indicated complete absence of the characteristic, 1 indicated moderate presence, and 2 indicated severe presentation. The rat pain grimace score (PGS) was calculated by averaging the scores given for each parameter for each photo, and averaging the score for a single animal over the specified time period^31, 32^.

### Statistical analysis

Statistical analyses were performed using Prism 7 (GraphPad). Power analysis was performed based on previous reports of changes in gut hypersensitivity in IBD models in rodents^33^ to predetermine sample size at a minimum of n = 5 rats in each experiment group. Rats were randomly allocated to groups in all experiments. The normality of data distribution was assessed using the Shapiro-Wilk test. In the experiments described in Figure 2, data were not normally distributed, and in these experiments, non-parametric tests were used. The Kruskal-Wallis test with post hoc Dunn’s test was used to compare more than two independent samples. The Mann-Whitney test was used for comparing two independent samples, and the Wilcoxon matched-pairs signed-rank test was used for paired values. For the normally distributed data, unpaired *t*-test, paired *t*-test, and ordinary and RM two-way ANOVA were used when appropriate. Actual p values are presented for each data set. The criterion for statistical significance was p < 0.05. Boxplots presented in all figures depict the mean (*solid line*), 25^th^, 75^th^ percentile, and 1.5 SD. Boxplots of not normally distributed data (Figure 2) also depict the median (*dotted line*). In some experiments (PGS, Figure 3), both males and females were used. To assess the presence of the main effects of sex or interactions of sex with experimental manipulations, these data sets were analyzed using sex as a factor. In no cases were any main effects of sex or interactions observed, so data were collapsed. In some experiments, only male rats were used.

**Figure 2.**
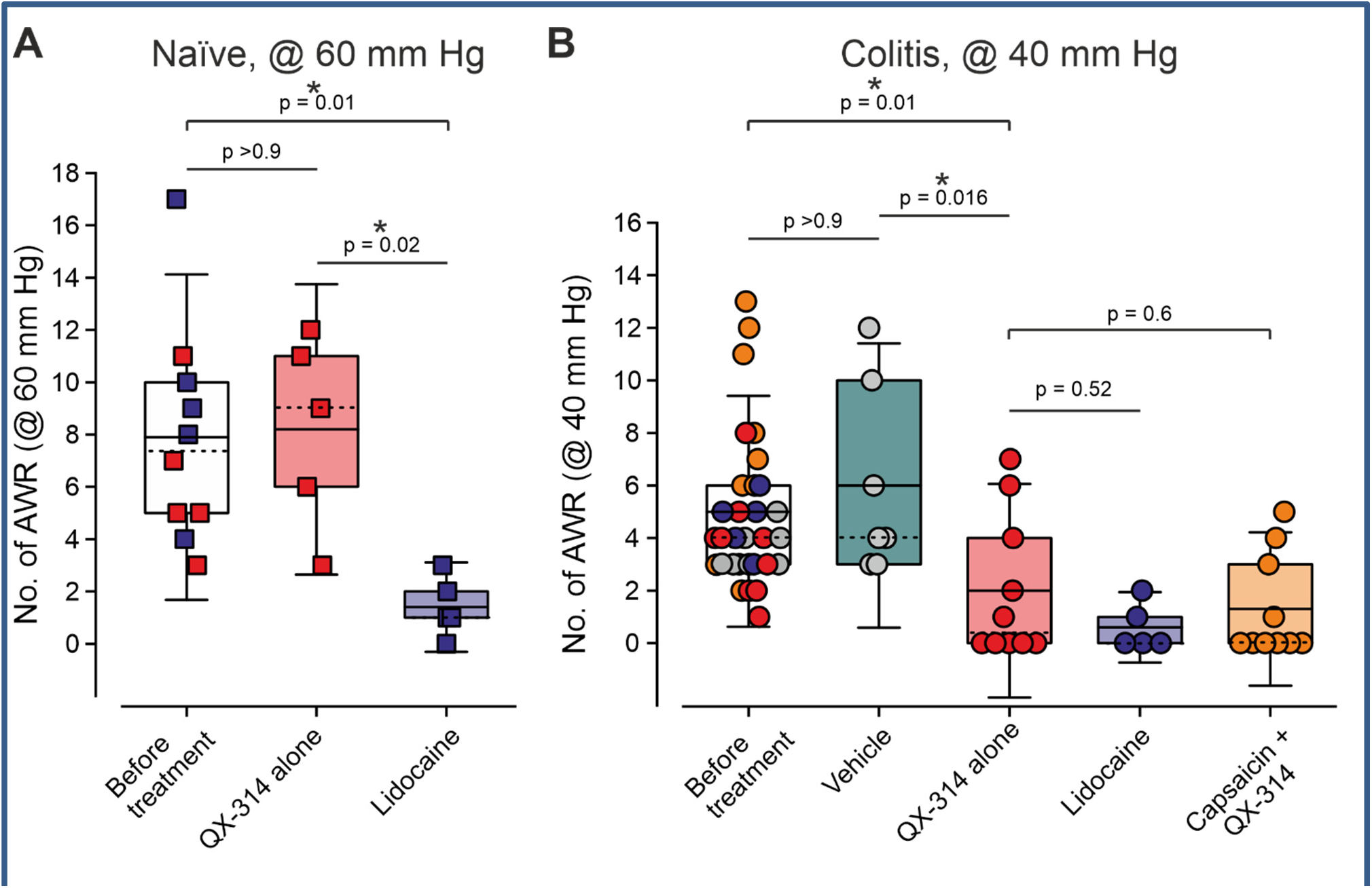
Application of QX-314 alone significantly reduces colitis-induced gut hypersensitivity but does not affect gut sensitivity in naïve rats. A. Box plot and individual values depicting the changes in the number of AWRs elicited in individual rats by colorectal distentions at 60 mm Hg, before and after treatment with 2% QX-314 alone and 2% lidocaine. Note that in these conditions, QX-314 does not affect gut sensitivity, and lidocaine significantly inhibits the response to colorectal distention. No difference in the number of AWRs before the treatment in the two groups (“QX-314” and “lidocaine”) was observed (p=0.28, Mann-Whitney test, N=5 rats in each group). In order to directly compare the effects of QX-314 and lidocaine in gut hypersensitivity, the values of the “Before” groups of the two treatments were combined, and the comparison was made between this group (N=10 rats) and “QX-314” and “lidocaine” groups using the Kruskal-Wallis test with post hoc Dunn’s test. The animals are color-coded according to treatment (QX-314 or lidocaine) groups. Importantly, the paired analysis of each treatment group showed similar significance (the comparison between the “Before” and “QX-314” groups (*red squares)*: p=0.12, N=5 rats; the comparison between “Before” and “Lidocaine” groups (*blue squares*): p=0.01, N=5 rats, Wilcoxon matched-pairs signed rank test). **B.** Same as A, but with rats in colitis condition. The AWRs were accessed following colorectal distention at 40 mm Hg and compared before and after treatment with vehicle, QX-314 alone, lidocaine, and capsaicin and QX-314 co-applied. Note that in these conditions, 2% QX-314 alone is sufficient to significantly reduce gut hypersensitivity. Similar to *A*, no difference in the number of AWRs before various treatments was observed; therefore, these values were combined (p=0.1, Kruskal-Wallis test with post hoc Dunn’s test, N=10 rats in “QX-314” group; N=7 in the “vehicle” group, N=5 in the “Lidocaine” group). In order to directly compare the effects of various treatments, the values assessed before the treatment in all groups were combined. For the “Capsaicin and QX-314” group, the data presented in Figure 1 was used (N=9 rats). Further comparison was made between this group (N=31 rats) and the effects of various treatments using the Kruskal-Wallis test with post hoc Dunn’s test. The animals are color-coded according to the treatment groups. Importantly, the paired analysis of each treatment to the values before the treatment showed similar significance (the comparison between the “Before” and “QX-314” groups (*red circles)*: p=0.03, Wilcoxon matched-pairs signed rank test, N=10 rats; the comparison between “Before” and “vehicle” groups (*gray circles*): p=0.18, Wilcoxon matched-pairs signed rank test, N=7 rats). The box plot depicts the mean (*solid line*), the median (*dotted line*), 25-75% percentile, and the whiskers depict 1.5 SD.

**Figure 3.**
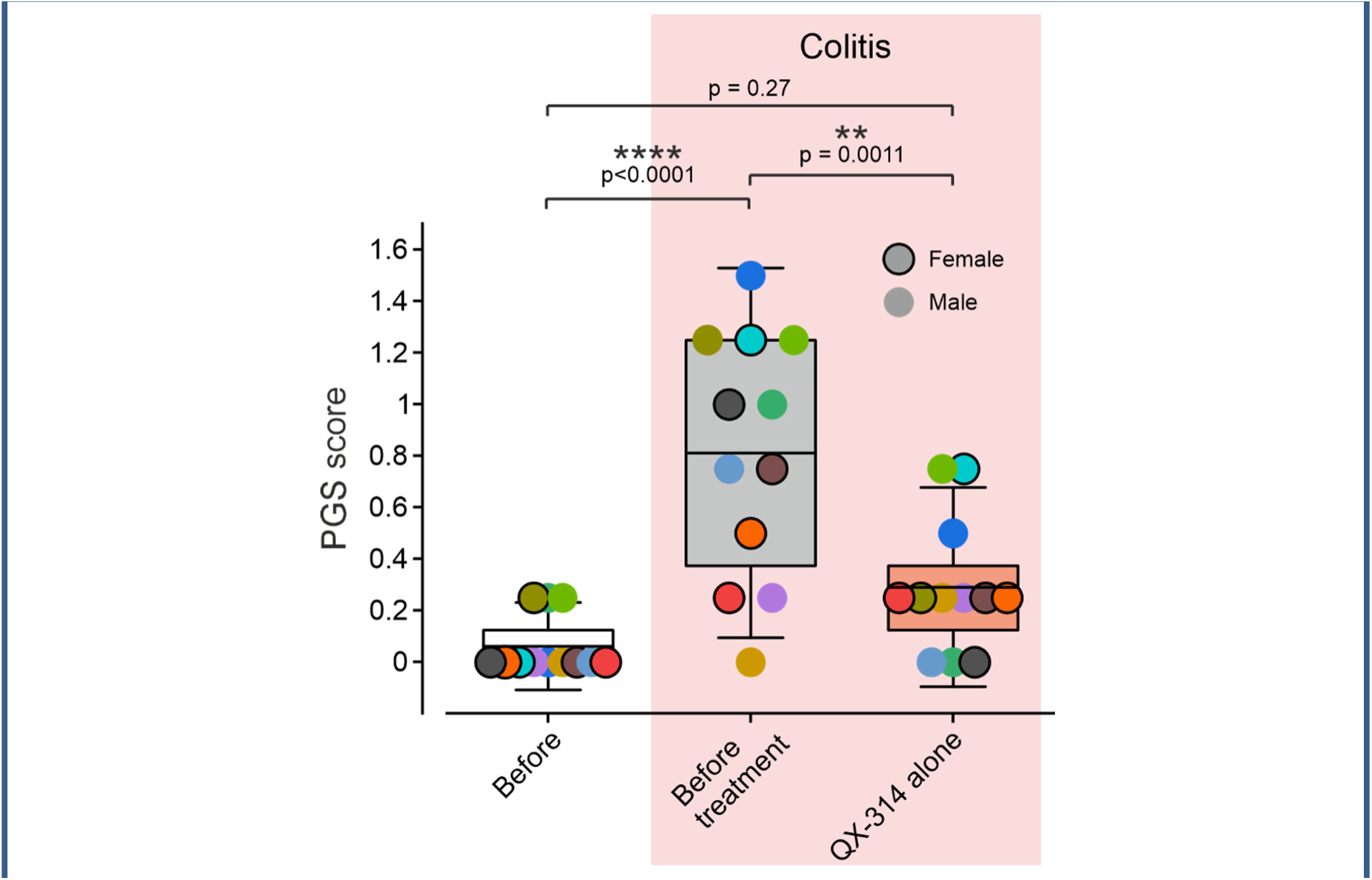
QX-314 alone significantly attenuates colitis-induced ongoing pain. A box plot and individual values of pain grimace scale (PGS) score assessed in the same rats (color-coded) before induction of colitis (“Before”) and 4 days after induction of colitis, before, and 10 minutes after intrarectal application of 2% QX-314 alone. RM one-way ANOVA with post hoc Bonferroni. N = 6 males and 6 female rats. The box plot depicts the mean, 25-75% percentile, and the whiskers depict 1.5 SD.

### Data availability

All datasets generated and/or analyzed during the current study are available in the main text or upon request. Further information and requests for resources and reagents should be directed to and will be fulfilled by the Lead Contact, Alexander Binshtok.

## Results

### Co-application of capsaicin with QX-314 into an inflamed colon diminishes colitis-induced visceral hypersensitivity

We examined the role of TRPV1-expressing nociceptive neurons in colitis-induced visceral hypersensitivity by monitoring the changes in visceral responsiveness to pressure following the silencing of these neurons. We used the abdominal withdrawal reflex (AWR) to estimate visceral hypersensitivity^33^ (**Figure 1A**). We measured the number of AWRs evoked in various intracolonic pressures, in naïve conditions, and in DNBS-induced colitis conditions (4 days after treatment with DNBS)^34^. In naïve animals, we did not detect any response when applying intracolonic pressure of 40 mm Hg (**Figure 1B**, *before,* see Methods). Applying the same pressure to the animals 4 days after DNBS led to substantial gut hypersensitivity (**Figure 1B, “***before treatment”*). All animals with increased gut sensitivity also showed prominent colonic inflammation **(Supplementary Figure 1**, *see Methods*). In these conditions, intrarectal application of capsaicin together with QX-314 produced a significant decrease in visceral hypersensitivity (**Figure 1B**) such that it reversed to the levels measured before inducing colitis (**Figure 1B**, *“before”*). Application of capsaicin alone to the inflamed gut led to substantial sensitivity to colorectal distention at 40 mm Hg; therefore, these experiments were terminated (*not shown*). Application of the vehicle (*see Methods*) did not affect colitis-induced visceral hypersensitivity (*see* **Figure 2B**). These results show that silencing of TRPV1-expressing neurons inhibits colitis-induced visceral hypersensitivity, suggesting the prominent role of these fibers in visceral pain.

### Application of QX-314 alone significantly affects visceral sensitivity in inflamed but not in naïve rats

Because QX-314 is a permanently charged compound, its penetration through the cell membrane is limited^35^. We and others have previously demonstrated that injection of QX-314 alone, either subcutaneously or peri-sciatically to naïve animals, in doses up to 2%, does not produce analgesia ^24, 36, 37^. Similarly, intrarectal application of 2% QX-314 alone to naïve animals did not change their responsiveness to intracolonic pressure, whereas application of 2% lidocaine fully inhibited it (**Figure 2A**). Notably, to evoke AWR in naïve animals, we used a higher pressure of 60 mmHg because we showed that 40 mmHg was insufficient to induce AWRs in naïve rats (see **Figure 1**). Surprisingly, applying 2% QX-314 alone in colitis conditions significantly reduced gut sensitivity (**Figure 2B**). The QX-314-induced analgesia was similar to the one produced by applying 2% lidocaine, or 2% QX-314 applied together with capsaicin (**Figure 2B**). using the Kruskal-Wallis test with post hoc Dunn’s test. The animals are color-coded according to treatment (QX-314 or lidocaine) groups. Importantly, the paired analysis of each treatment group showed similar significance (the comparison between the “Before” and “QX-314” groups (*red squares)*: p=0.12, N=5 rats; the comparison between “Before” and “Lidocaine” groups (*blue squares*): p=0.01, N=5 rats, Wilcoxon matched-pairs signed rank test). **B.** Same as A, but with rats in colitis condition. The AWRs were accessed following colorectal distention at 40 mm Hg and compared before and after treatment with vehicle, QX-314 alone, lidocaine, and capsaicin and QX-314 co-applied. Note that in these conditions, 2% QX-314 alone is sufficient to significantly reduce gut hypersensitivity. Similar to *A*, no difference in the number of AWRs before various treatments was observed; therefore, these values were combined (p=0.1, Kruskal-Wallis test with post hoc Dunn’s test, N=10 rats in “QX-314” group; N=7 in the “vehicle” group, N=5 in the “Lidocaine” group). In order to directly compare the effects of various treatments, the values assessed before the treatment in all groups were combined. For the “Capsaicin and QX-314” group, the data presented in Figure 1 was used (N=9 rats). Further comparison was made between this group (N=31 rats) and the effects of various treatments using the Kruskal-Wallis test with post hoc Dunn’s test. The animals are color-coded according to the treatment groups. Importantly, the paired analysis of each treatment to the values before the treatment showed similar significance (the comparison between the “Before” and “QX-314” groups (*red circles)*: p=0.03, Wilcoxon matched-pairs signed rank test, N=10 rats; the comparison between “Before” and “vehicle” groups (*gray circles*): p=0.18, Wilcoxon matched-pairs signed rank test, N=7 rats). The box plot depicts the mean (*solid line*), the median (*dotted line*), 25-75% percentile, and the whiskers depict 1.5 SD.

### Application of QX-314 alone significantly reduces colitis-induced non-reflexive, ongoing pain

So far, we have assessed colitis-induced pain by measuring changes in AWR evoked by applying visceral distension. This widely accepted approach provides a good estimation of changes in evoked visceral sensitivity in various conditions, for example^9, 38^. Another important and clinically relevant question is whether DNBS-induced colitis leads to non-evoked ongoing pain and the effect of selective silencing of visceral nociceptors on ongoing pain. To answer these questions, we assessed the levels of colitis-induced ongoing pain by evaluating a rat pain grimace scale (PGS)^31, 32^. As expected, before inducing colitis, rats did not exhibit facial characteristics of ongoing pain (**Figure 3**, **Supplementary Figure 2A**). Four days after inducing colitis, the PGS score was significantly increased, as rats exhibited substantial orbital tightening, cheek bulging, ear flattening, and whisker stiffening (**Supplementary Figure 2B**), the markers of spontaneous pain^31, 32^. Notably, the application of 2% QX-314 alone alleviated the signs of spontaneous pain, reversing the PGS score to the values measured before colitis induction (**Figure 3, Supplementary Figure 2C**).

Altogether, our results show that applying QX-314 alone is sufficient to reduce both colitis-induced evoked visceral hypersensitivity and ongoing pain. These results imply that in the inflamed gut, QX-314 does not require exogenic activation of TRPV1 channels to access the cytoplasm of nociceptive neurons, suggesting that these channels are tonically activated due to inflammation.

### Local blockade of TRPV1 channels prevents the analgesic effect of QX-314

What could explain the analgesic effect of membrane impermeant QX-314 in an inflamed colon? DNBS-induced gut inflammation could potentially lead to an increase in mucosal and blood vessel permeability, allowing QX-314 to reach the blood flow and affect pain systemically. If the latter is the case, QX-314 applied to the inflamed guts should have a general inhibitory effect on the responses to noxious stimuli. However, this is less likely considering the low membrane permeability of QX-314. Indeed, in the colitis conditions, we did not detect any effect of QX-314 applied intrarectally on the thermal pain threshold measured from the rats’ paws (**Figure 4**).

**Figure 4.**
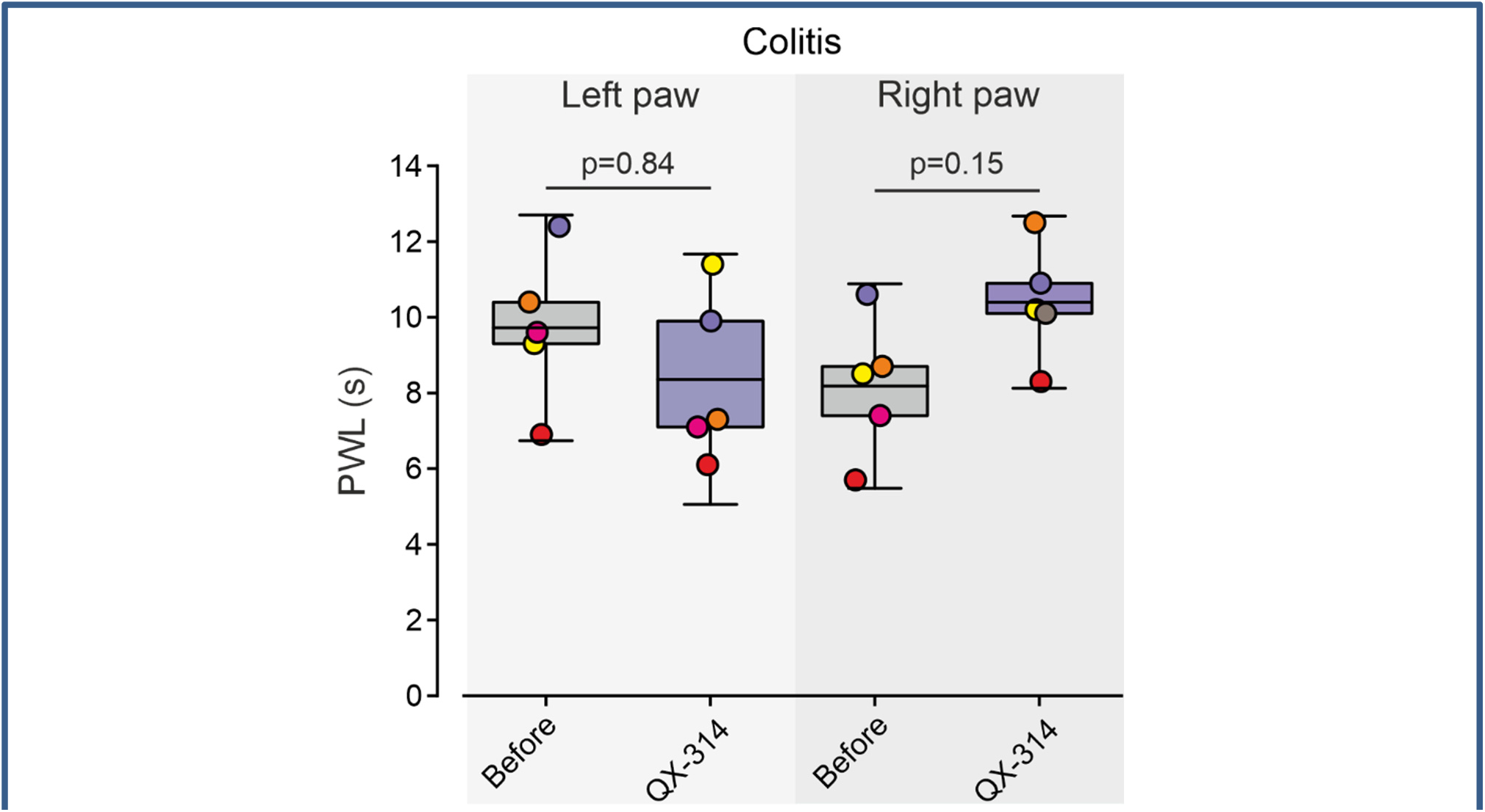
QX-314 applied to the colitic colon does not have a systemic effect on sensitivity to noxious stimuli. A box plot and individual values of the latency to the withdrawal response to a noxious heat stimuli (paw withdrawal latency, PWL) applied to the left and right hind paw of the same rats (color-coded) in colitis conditions, assessed before, and 10 minutes after intrarectal application of QX-314. Note, no effect of QX-314 on the responsiveness to the noxious stimuli applied to the paw. Paired *t*-test. N = 5 rats. The box plot depicts the mean, 25-75% percentile, and the whiskers depict 1.5 SD.

Alternatively, QX-314 could enter nociceptive neurons innervating the gut via inflammation-mediated tonically active TRPV1 channels. The role of tonically active TRPV1 channels in inflammatory conditions, but not in IBD models, was previously suggested^26, 39^. Moreover, it has been shown that this activation of TRPV1 channels is sufficient to allow the entry of QX-314 into neurons leading to the inhibition of their activity^25, 26, 39^. To examine the hypothesis that in the colitis model also, QX-314 accesses nociceptive neurons via tonically active TRPV1 channels, we pharmacologically inhibited TRPV1 channels and studied the effect of QX-314 on gut sensitivity in these conditions. To that end, we used TRPV1 selective antagonist capsazepine^40^. Because capsazepine was not used before to inhibit colon-rectal TRPV1 channels, we first determined the proper dose of capsazepine, sufficient to inhibit gut TRPV1 channel activation **(Figure 5A**). We administered naïve rats with subsequently increasing doses of intrarectal capsazepine followed by an application of 1 mg/ml capsaicin, which, when applied alone, produced a substantial increase in gut sensitivity **(Figure 5A**). We chose the dose of capsazepine, which fully prevented capsaicin-induced gut hypersensitivity (3 mg/ml, **Figure 5A**), and applied capsazepine to the DNBS-treated animals (**Figure 5B**). In these conditions, the co-application of capsazepine and QX-314 prevented the analgesic effect of QX-314 (**Figure 5B**), which we previously described (see **Figure 2B**). The application of capsazepine alone did not affect colitis-induced visceral hypersensitivity (**Figure 5B**).

**Figure 5.**
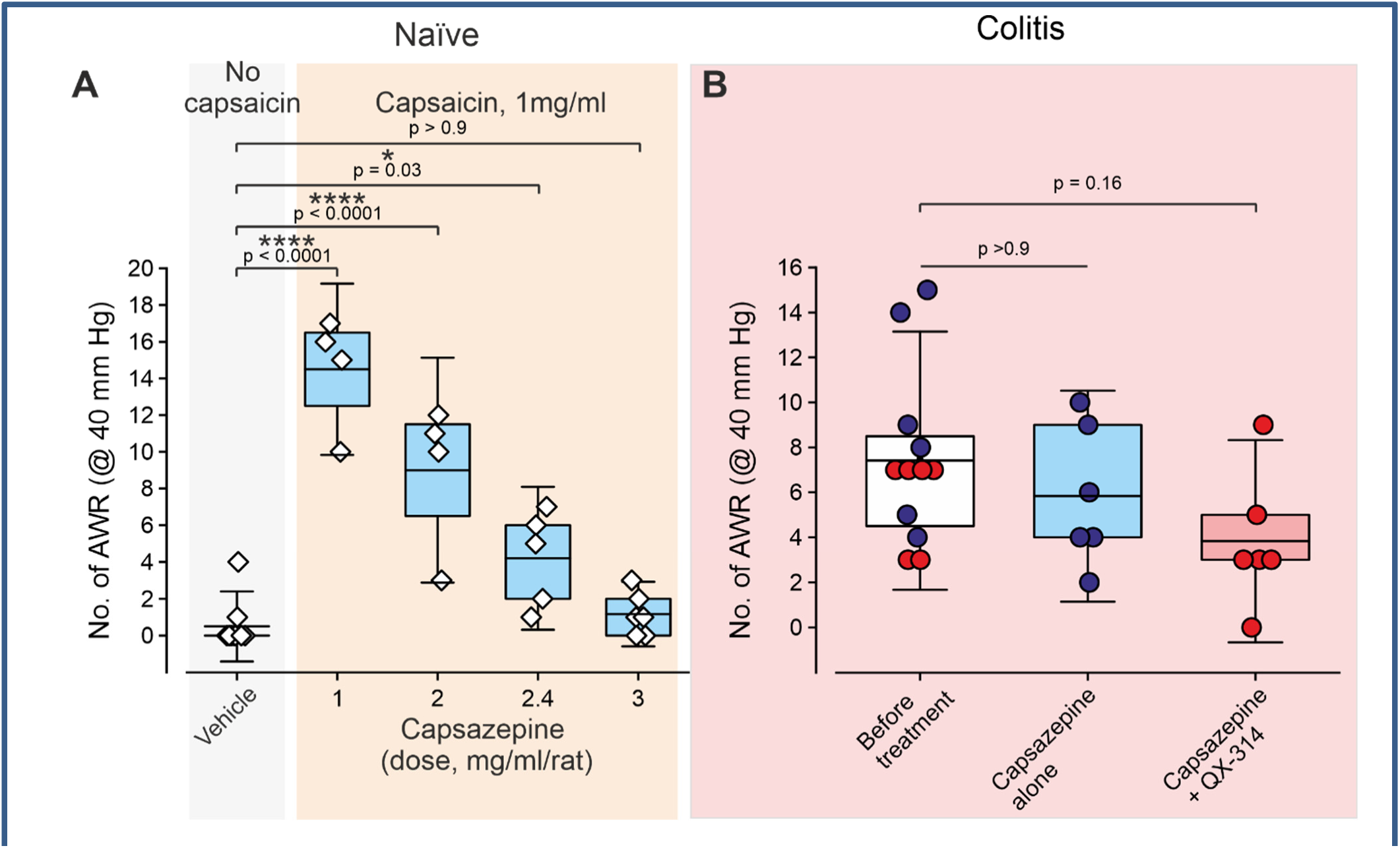
Inhibition of TRPV1 channels prevents the analgesic effect of QX-314 in gut hypersensitivity. A. Box plot and individual values depicting the changes in the number of AWRs elicited in individual rats by colorectal distentions at 40 mm Hg following application of vehicle and capsaicin together with capsazepine at the indicated doses. Note that 3 mg/ml of capsazepine prevented the irritant effect of capsaicin, and this dose was used in the following experiments. Ordinary one-way ANOVA with post hoc Bonferroni. **B.** Box plot and individual values depicting the changes in the number of AWRs elicited in individual rats, 4 days after induction of colitis, by colorectal distentions at 40 mm Hg, before and after treatment with capsazepine alone, or capsazepine with QX-314. Note that in colitis conditions, QX-314 applied with capsazepine does not affect gut hypersensitivity. No difference in the number of AWRs before treatments with capsazepine alone or capsazepine together with QX-314 was observed (p=0.08, unpaired *t*-test, N=6 rats in each group); thus, the values of the AWRs of the groups before applying the treatment groups were pooled. Further comparison was made between this group (N=12 rats) and treatment groups using the ordinary one-way ANOVA with post hoc Bonferroni. The animals are color-coded according to the treatment groups. Importantly, the paired analysis of the effect of treatment of each group separately showed similar significance (the comparison between the “Before” and “Capsazepine alone” groups (*red squares*): p=0.1, N=6 rats; the comparison between “Before” and “Lidocaine” groups (*blue squares*): p=0.12, N=6 rats, paired *t*-test). The box plot depicts the mean, 25-75% percentile, and the whiskers depict 1.5 SD.

These data imply that TRPV1 channels are tonically active in DNBS-induced colitis and suggest that QX-314 affects gut hypersensitivity by entering nociceptive neurons, at least in part, via these tonically active TRPV1 channels.

## Discussion

Despite the plethora of anti-inflammatory drugs available or in the pipeline for the treatment of IBD-associated inflammation, there is an unmet need for treating IBD-associated abdominal pain. In the current paper, we advance the current understanding of the molecular and cellular pathways involved in abdominal pain, in a rodent model of IBD, with potential implications for drug development. We show that local administration of a membrane impermeant sodium channel blocker QX-314 into the inflamed colon of a rat caused a decrease in visceral-specific hypersensitivity and also decreased ongoing pain behavior. We further show that this analgesic effect was unique to colitis and was not demonstrated in naïve animals. Furthermore, we show that co-administering blockers of the TRPV1 channel prevented this effect. Taken together, our findings provide novel evidence proposing that TRPV1 channels in the inflamed gut may be tonically active, allowing the entry of charged and, therefore, membrane impermeant analgesic molecules selectively into nociceptive fibers.

TRP channels, mainly TRPV1 and TRPA1, have been extensively studied as both culprits and possible therapeutic targets in IBD-associated pain and inflammation^12, 41^. These channels are mainly expressed on the extrinsic nociceptive neuronal ending in the gut, but recent evidence suggests they can also be found on intrinsic sensory neurons, surface epithelial cells and inflammatory cells, therefore providing a possible link between sensory pathway activation and neurogenic inflammation^11, 12^. Indeed, it was suggested that blockade or genetic deletion of these channels in rodent IBD models may affect gut inflammation and thereby reduce visceral pain^42, 43^. Unfortunately, no current therapeutic option exists for the selective blockade of TRPV1 channels, as the available TRPV1 antagonists cause substantial side effects in humans. Selective silencing of TRPV1-expressing visceral nociceptors, which we report here, in addition to its direct effect on pain, could also, as previously reported, diminish neurogenic inflammation^26, 39^, which could further decrease inflammatory burden in IBD.

Counter-intuitively, the TRPV1 antagonist capsazepine used in our study did not significantly affect visceral hypersensitivity in the inflammatory state. These findings contrast previous reports^44–46^. The difference in the delivery method of the drug could explain this discrepancy. In previous publications, TRPV1 antagonists were delivered intraperitoneal or intrathecally, likely acting at multiple levels of pain transduction. We are the first to report on the use of locally administered intra-colonic capsazepine, requiring us to perform preliminary dose-finding experiments in naive animals. It is possible that applying a higher dose of capsazepine would lead to a complete blockade of TRPV1 channels and thus affect, at least in part, visceral hypersensitivity.

Our study suggests that tonic activation of TRPV1 channels contributes to IBD pain. Previous studies suggest that abnormal activity of other channels, such as TRPA1^47^ or Piezo 2^8, 10^ also plays a role in the pathophysiology of IBD pain. Importantly, we and others demonstrated that abnormal activity of TRPA1 can also provide an entry of QX-314 into neurons^25, 48^. Nevertheless, the fact that these channels are expressed on TRPV1-containing neurons^10, 49^ suggests that our selective silencing approach would still block the activity of these neurons regardless of its molecular origin. While in this study, we used QX-314 as a proof-of-concept for selective silencing of visceral nociceptors, the use of QX-314 by humans is limited by its neurotoxicity ^50, 51^, and, therefore, not viable for a translational approach. Newly developed charged ion channel blockers^39^ may prove to be safer to use in humans, especially if delivered locally to the gut. Further challenges will be to ensure a long-acting effect and to fine-tune these platforms to maximize their effectiveness.

In summary, our results suggest the central role of TRPV1-expressing nociceptive neurons and tonic activation of TRPV1 channels in IBD pain. Moreover, our findings provide a conceptual approach for selective inhibition of IBD pain by buttressing a basis for employing charged activity blockers to selectively and effectively block visceral pain.

## Authors contributions

Conceptualization, Y.M. and A.M.B.; Methodology Y.M., N.E., S.L., H.N., L.R. and A.M.B.; Investigation, Y.M., N.E., S.L., H.N., L.R. and A.M.B.; Writing – Original Draft, Y.M., N.E., S.L., and A.M.B.; Funding Acquisition, A.M.B.; Supervision, A.M.B.

## Acknowledgments

*Funding sources*: Support is gratefully acknowledged from the Israeli Cancer Research Foundation (ICRF, Brause Initiative Application), Canadian Institute of Health Research (CIHR), the International Development Research Centre (IDRC), the Israel Science Foundation (ISF) and the Azrieli Foundation – grant agreement 2545/18; Israeli Science Foundation – grant agreement 1470/17; the Deutsch-Israelische Projectkooperation program of the Deutsche Forschungsgemeinschaft (DIP) grant agreement B.I. 1665/1-1ZI1172/12-1 and Sessile and Seymour Alpert Chair in Pain Research.

