## Supplementary Figures and legends for "Attenuation of colitis-induced visceral hypersensitivity and pain by silencing TRPV1-expressing fibers in rat colon"

**Supplementary Figure 1**

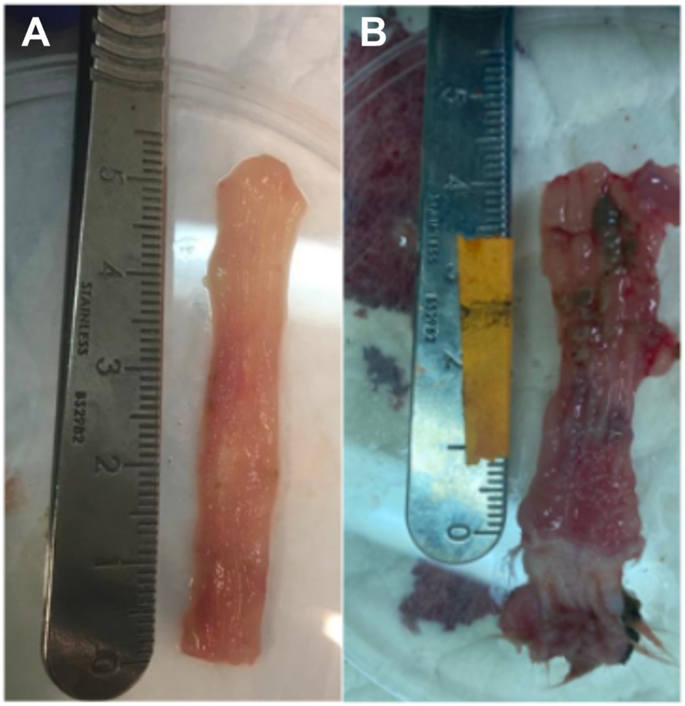


**Supplementary Figure 1.** Ano-colon, resected and dissected open from naïve rats (A) and rats at day 4 following DNBS-induced colitis (B). In panel B, inflammation involving up to 4 cm of the distal colon can be seen, with variable areas of erythema, mucosal fold thickening, creeping fat, and focal necrosis.

**Supplementary Figure 2**

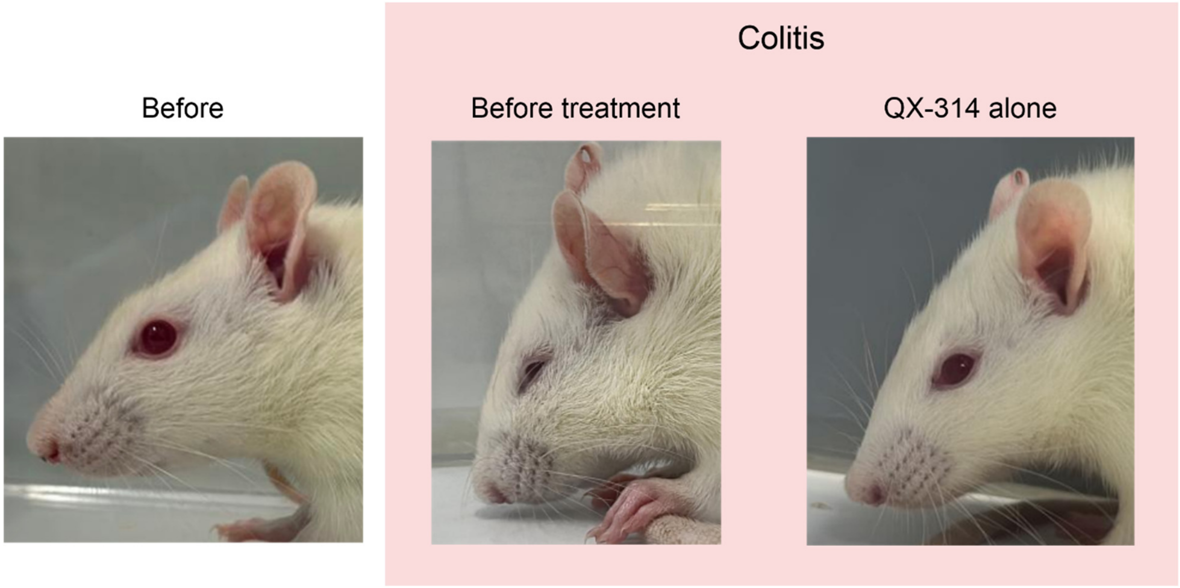


**Supplementary Figure 2.** Representative images of a face of an individual male rat (represented as *blue* in Figure 3) were taken before and after inducing colitis and 10 minutes after treatment with QX-314. Note that after inducing colitis rat exhibits prominent orbital tightening, cheek bulging, ear flattening, and whisker stiffening and that these signs disappear after treatment with QX-314 alone.
